# Evolution of an alternative genetic code in the *Providencia* symbiont of the haematophagous leech *Haementeria acuecueyetzin*

**DOI:** 10.1101/2023.03.27.534433

**Authors:** Alejandro Manzano-Marín, Sebastian Kvist, Alejandro Oceguera-Figueroa

## Abstract

Strict blood-feeding animals are confronted with a strong B vitamin deficiency. Blood-feeding leeches from the Glossiphoniidae family, similarly to haematophagous insects, have evolved specialised organs called bacteriomes to harbour symbiotic bacteria. Leeches of the *Haementeria* genus have two pairs of globular bacteriomes attached to the oesophagus which house intracellular ’*Candidatus* Providencia siddallii’ bacteria. Previous work analysing a draft genome of the *Providencia* symbiont of the Mexican leech *Haementeria officinalis* showed that, in this species, the bacteria hold a reduced genome capable of synthesising B vitamins. In this work, we aimed to expand our knowledge on the diversity and evolution of *Providencia* symbionts of *Haementeria*. For this purpose, we sequenced the symbiont genomes of three selected leech species. We found that all genomes are highly syntenic and have kept a stable genetic repertoire, mirroring ancient insect endosymbionts. Additionally, we found B vitamin pathways to be conserved among these symbionts, pointing to a conserved symbiotic role. Lastly and most notably, we found that the symbiont of *Haementeria acuecueyetzin* has evolved an alternative genetic code, affecting a portion of its proteome and showing evidence of a lineage-specific and likely intermediate stage of genetic code reassignment.

## Introduction

Obligate nutritional symbioses are widespread in animals with a restricted diet, and have been most extensively studied in phloem-feeding arthropods. Numerous lineages of these symbiotic organisms have convergently evolved characteristic genomic features, including a reduced gene-dense genome, a biased G+C content, a lack of mobile genetic elements, and elevated substitution rates particularly at sites affecting amino acid substitutions (Baumann, 2005; Latorre and Manzano-Marín, 2017; McCutcheon and Moran, 2012; McCutcheon *et al*., 2019; Moran *et al*., 2008). The study of these nutritional symbionts has mostly focused on plant-feeding arthropods, while historically largely overlooking invertebrates feeding on other nutrient-deficient diets. One such group is that of strict blood-feeders, where their diet is distinguished by a strong B-vitamin deficiency (Lehane, 2005). In blood-feeding arthropods, bacteria housed in so-called bacteriomes (specialised organs evolved to house symbiotic bacteria) compensate for this deficiency, by producing and delivering these nutrients to the host (Kir, 2010; Duron *et al*., 2018; Nikoh *et al*., 2014; Nogge, 1981; Nogge and Gerresheim, 1982). Leeches (Annelida: Hirudinida) are a monophyletic group of annelids that are most famous for their widespread blood-feeding habit (Tessler *et al*., 2018), Within the family Glossiphoniidae, the ancestral evolution of a proboscis (Trontelj *et al*., 1999), a tubular mouthpart used to penetrate host tissues, results in members of this family being able to feed namely on a liquid or soft-tissue based diet, consisting of blood or invertebrate haemolymph (Sawyer, 1986). Similarly to strict blood-feeding arthropods, species of the *Haementeria, Placobdella*, and *Placobdelloides* genera possess distinctive organs attached to the oesophagus in which symbiotic bacteria reside (Kikuchi and Fukatsu, 2002; Perkins *et al*., 2005; Siddall *et al*., 2004). In the case of *Haementeria* leeches, two distinct pairs of globular sacs (also known as bacteriomes) are attached to the oesophagus by thin ducts (Oceguera-Figueroa, 2006, 2008), and microscopic investigations have corroborated the presence of such bacteria in the bacteriomes of species in this genus (Manzano-Marín *et al*., 2015; Perkins *et al*., 2005).

*Haementeria* is a strictly New World leech genus with two main lineages: a Central and South American clade (H. *tuberculifera, H. lutzi*, and H. *paraguayensis*), and a Mexican and South American clade made up of the species *H. acuecueyetzin, H. officinalis, H. depressa, H. ghilianii*, and *H. lopezi* (**figure 1**) (Oceguera-Figueroa, 2012). All species form the latter clade have been identified to carry and have co-diverged with the obligate symbiont *P. siddallii* (Manzano-Marín *et al*., 2015). Previous genomic sequencing of the bacteria housed in the bacteriomes of the Mexican leech *Haementeria officilalis* revealed a singular bacterial species ’*Candidatus* Providencia siddallii’ (hereafter referred to as *P. siddallii*) inhabiting the cytoplasm of bacteriocytes (Manzano-Marín *et al*., 2015). Genome-based metabolic analysis revealed a large portion of its genetic repertoire is dedicated to fulfilling its symbiotic role: biosynthesising B vitamins to supplement the host’s deficient blood-based diet. This study revealed a convergent retention of the B-vitamin biosynthetic pathways among endosymbionts of distantly related blood-feeding insect taxa and the leech *H. officinalis*. Mirroring what has been observed for numerous strict vertically-transmitted obligate nutritional endosymbionts in sap- and blood-feeding insects, the genome of *P. siddallii* evidenced a historical past involving large-scale gene loss and genomic deletions (Manzano-Marín *et al*., 2015). This genomic reduction process is due to a combination of relaxed selection on dispensable genes and the strong bottlenecks resulting from the strict vertical transmission maternally-inherited endosymbionts go through in each generation, increasing the fixation of deleterious mutations through drift (Bennett and Moran, 2015; Moran, 1996). In its most advanced stages, this extreme evolutionary process can lead to an adapt-or-die scenario, where mutations arising in essential genes can lead to the breakdown of the symbiosis (resulting in the extinction of the host lineage or symbiont replacement) or to compensatory mutations in one or both partners (Bennett *et al*., 2016; Latorre and Manzano-Marín, 2017; McCutcheon and Moran, 2012; McCutcheon *et al*., 2019).

**Figure 1.**
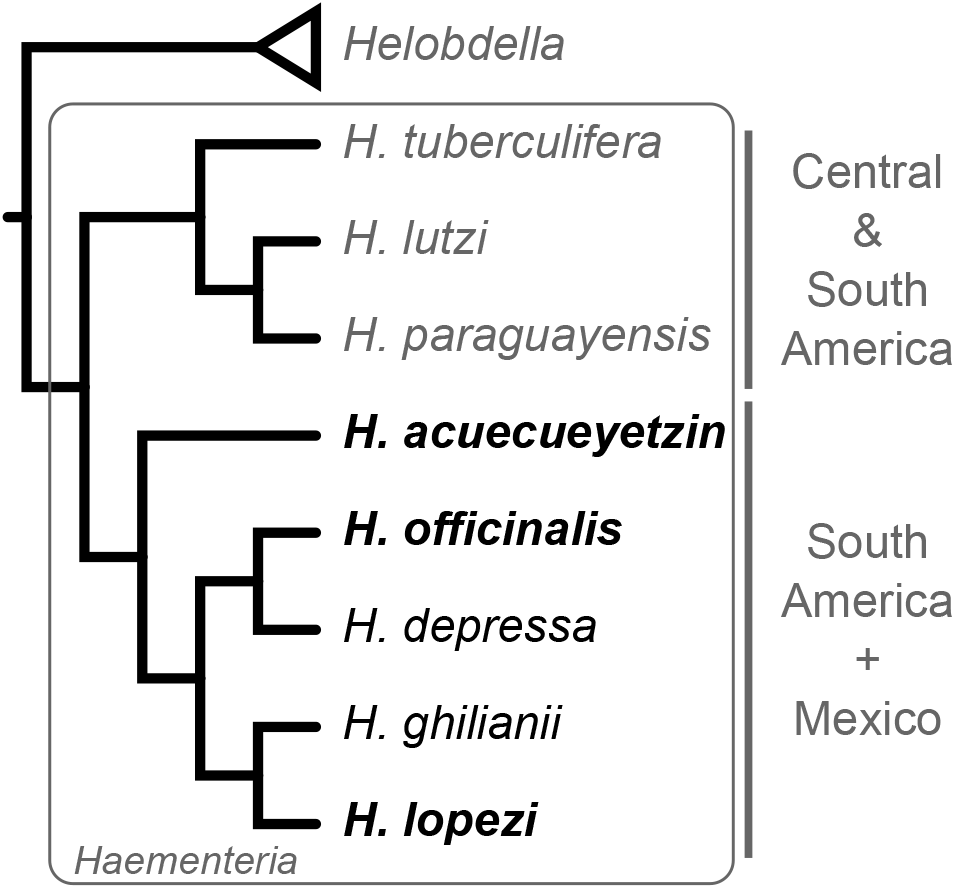
Phylogenetic relationships of *Haementeria* species. Dendrogram displaying the phylogenetic relationships among *Haementeria* species according to Oceguera-Figueroa (2012). Mexican species whose endosymbionts were sequenced in this study are highlighted in bold.

In this work, we aimed to gain deeper insights into the *Haementeria*-*Providencia* symbiosis. For this purpose, we sequenced the genome of the *P. siddallii* endosymbionts of the Mexican leeches *H. acuecueyetzin* and *H. lopezi*. In addition, we have re-sequenced and produced a closed reference genome for the previously-sequenced *P. siddallii* endosymbiont of *H. officinalis*. It is worth noting that Mexican species of *Hamenteria* do not form a monophyletic group. On the contrary, *H. lopezi* and *H. officinalis* are sister to their respective South American counterparts (*H. ghilianii* and *H. gracilis*), whereas *H. acuecueyetzin* represents an independent lineage (**figure 1**). Through comparative genomics, we have found *P. siddallii* symbionts display large-scale genome synteny, conservation of enzymes involved in B-vitamin provisioning, no mobile elements, and little variation in gene content. Most notably, we discovered a rare genetic code change from the traditional bacterial code 11 (identical to the standard, but including the alternative initiation codons AUU, AUC, and AUA) (Elzanowski and Ostell, 2019) to the rarer genetic code 4, where the UGA codon is translated to tryptophan instead of being interpreted as a STOP. Finally, through thorough analysis of the protein-coding and tRNA-Trp (*trnW*) genes of the *P. siddallii* genomes, we conclude that the symbiont of *H. acuecueyetzin* displays a lineage-specific, and likely intermediate, stage of the genetic code recoding process.

## Results

### *Providencia siddallii* genomes and biosynthesis of B vitamins

The genomes of the *P. siddallii* endosymbionts of *H. acuecueyetzin, H. officinalis*, and *H. lopezi* (hereafter **PSAC, PSOF** and **PSLP**, respectively) share many general genomic characteristics (**table 1**). All hold a compact genome of under 0.9 Mega base-pairs, a low G+C content, and a reduced set of genes, when compared to the free-living *Providencia rettgeri* strain. All three *P. siddallii* genomes also show a reduced set of non-coding RNAs, including only one ribosomal rRNA gene set, with separate 16S and 23S+5S rRNA genes. PSOF retains the largest genome, around 100 kilo base-pairs (**kbp**) larger than those of PSAC and PSLP, while retaining a similar amount of genes. A phylogenetic reconstruction using the conserved set of ribosomal proteins reiterated the co-divergence of *P. siddallii* and *Haementeria* (supplementary **figure S1**). It also revealed that the closest free-living relative of *P. siddallii* strains is *Providencia heimbachae*, that similarly to other *Providencia* strains, has been commonly isolated as an animal pathogen (O’Hara *et al*., 2000). In light of the co-divergence of *P. siddalllii* symbionts and their hosts, *P. siddalllii* has undergone at least two independent events of drastic genome reduction after the diversification of their hosts, which have almost exclusively impacted non-coding DNA. In addition, the genomes all show a small number of pseudogenes as well as a complete lack of any detectable mobile elements.

**Table 1.** General genomic characteristics of *P. siddallii* endosymbionts and the closely related free-living *Providencia rettgeri* strain AR 0082. The newly sequenced organisms are highlighted with a grey background.

| Bacterial strain | *P. rettgeri* AR_0082 | *P. siddallii* |  |  |
| --- | --- | --- | --- | --- |
|  |  | PSAC | PSOF | PSLP |
| genome size | 4,454 kbp | 756 kbp | 844 kbp | 741 kbp |
| G+C content | 40.3 % | 22.0 % | 23.9 % | 23.3% |
| CDSs | 4,071 | 610 | 628 | 612 |
| rRNAs | 22 | 3 | 3 | 3 |
| tRNAs | 77 | 33 | 33 | 34 |
| ncRNAs | 3 | 2 | 2 | 2 |
| pseudogenes | 74 | 4 | 5 | 6 |
| Mobile elements | 16 | None | None | None |

*P. siddallii* strains show large-scale genome synteny (supplementary **figure S2**) and share 96.27% of their protein-coding genes (**figure 2A**). The strain-specific gene content reflects mainly a differential retention of genes, mostly affecting cellular maintenance, regulation, and RNA modifications. Notably, PSOF and PSLP retain a slightly larger repertoire of proteases (*ptrB* and *pepQ* in both, *htpX* in PSOF), and genes involved in bacterial outer membrane biogenesis (*lpxL* and *hldD* in both, *mltB* and *gmhB* in PSOF) and cellular maintenance (*pcnB, holC, holD, ftsX*, and *ftsE* in both; and *rmuC* and *engB* in PSOF). As observed in other blood-feeder’s endosymbionts, PSAC and PSLP preserve intact and identical pathways to PSOF for the biosynthesis of B vitamins, putatively required to compensate for the host’s deficient diet (**figure 2B**; Manzano-Marín *et al*., 2015). All *Haementeria* symbionts have lost the *ilvD, ilvE*, and *panD* genes, rendering them able to synthesise pantothenate only from *α*-ketovaline and *β*-alanine. In addition, none retains a *nudB* gene, coding for a dihydroneopterin triphosphate diphosphatase. All genomes contain a small number of protein-coding genes (CDSs) with frameshifts in poly(A/T) tracts (supplementary **table S1, figure S3**) which resemble those reported to be rescued by transcriptional slippage in strains of *Buchnera aphidicola* and *Blochmannia pensilvanicus* endosymbionts (Tamas *et al*., 2008). From these, three genes containing such tracts have likely evolved these regions before the diversification of the symbiont lineage: *rlmE, murI*, and *znuA*, involved in 23S rRNA modification, cell wall biosynthesis, and zinc uptake, respectively.

**Figure 2.**
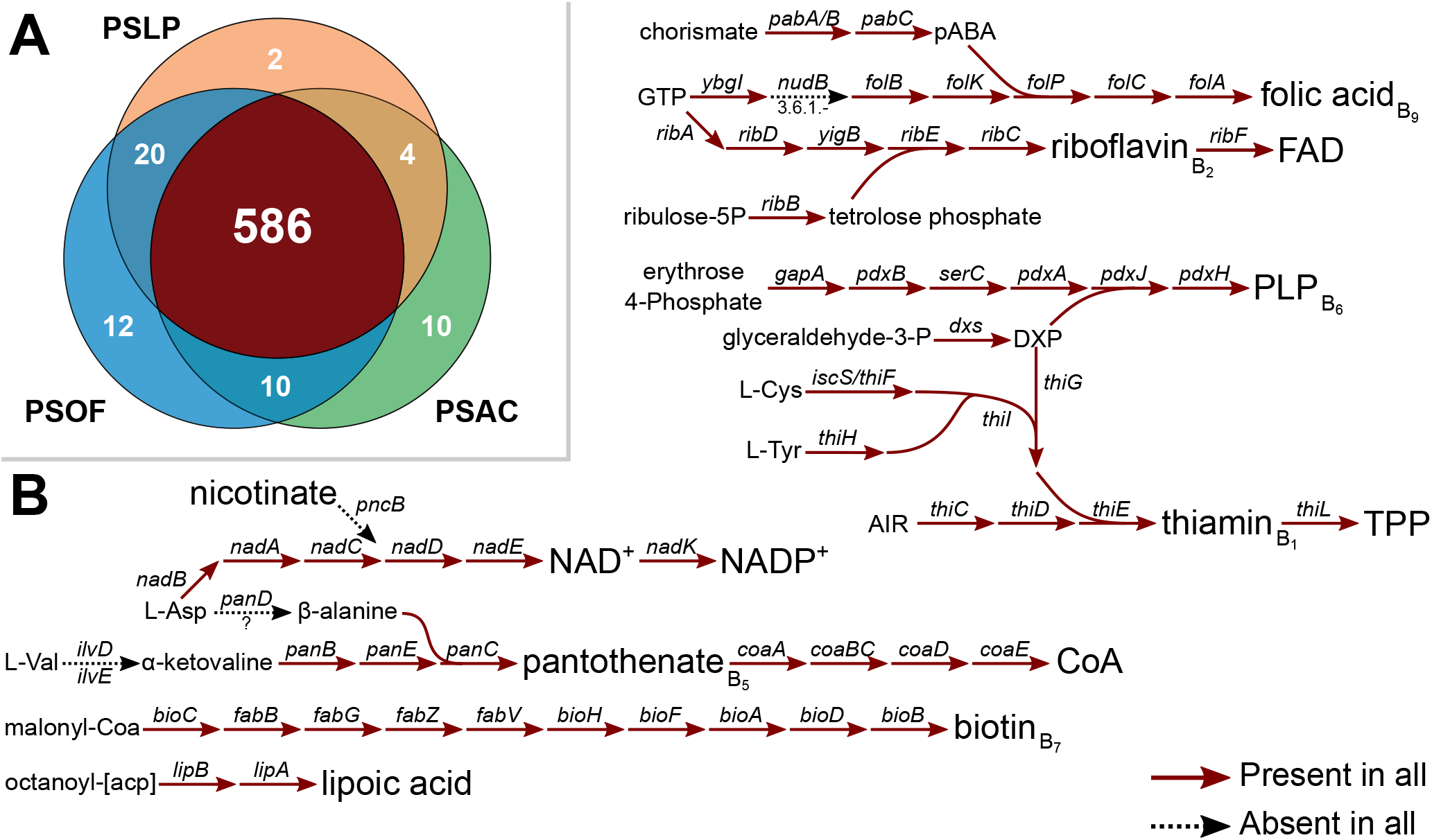
*P. siddallii* shared gene content and B vitamin biosynthesis. In the top left, Venn-like diagram displaying the results of OrthoMCL clustering of the predicted proteomes of PSAC, PSOF, and PSLP. In the rest of the figure, diagram of the B vitamin biosynthetic pathways. Arrows connect metabolites, and names on arrows indicate the gene name coding for the enzyme involved in the enzymatic step.

### Evolution of an alternative genetic code in the symbiont of *H. acuecueyetzin*

During the annotation process of PSAC, an unusually large number of proteins (125 out of 610) were found to be truncated by an early in-frame UG**A** stop codon. Upon closer inspection, most of these cases corresponded to UG**G**>UG**A** mutations (Tryptophan to STOP), as judged by comparing PSOF, PSLP, and other *Providencia* to PSAC (**figure 3A**). These UG**A**-containing genes include many whose products are considered essential for cellular maintenance as well as its symbiotic nutrient-provisioning role. These include genes coding for enzymes of the NADH-quinone oxidoreductase (*nuoC, nuoE, nuoG, nuoK*, and *nuoM*), aminoacyl--tRNA ligases (*leuS, glnS, ileS, gltX, cysS, glyQ, pheT*, and *valS*), biotin, thiamin, and folic acid biosynthesis (*bioA, thiE, thiF, thiH, pabC, folA, dxs, folP*, and *iscS*). This large quantity of putative UG**A**-containing genes suggested a genetic code change from 11 to 4, where the difference lies with the UG**A** codon coding for the amino acid tryptophan rather than for a stop codon recognised by the *prfB* gene product (peptide chain release factor 2, or **RF2**). This ”11-to-4” change has most famously evolved in *Entomoplasmatales* and *Mycoplasmatales* (*Mollicutes*), such as *Mycoplasma* spp. (Bové, 1993), and was first described in *M. capricolum* (Yamao, 1985). Beyond these, genetic code changes re-assigning STOP codons are known to have evolved in distantantly related lineages, such as in *Blastocrithidia* (Trypanosomatidae; Záhonová *et al*., 2016), ’*Candidatus* Absconditabacteria’ (or SR1 bacteria; Campbell *et al*., 2013), and some diplomonads (Keeling and Doolittle, 1996), among others (Ivanova *et al*., 2014; Shulgina and Eddy, 2021). To our knowledge, only four other such cases off ”11-to-4” change have been reported for nutritional/digestive endosymbionts: once in the alphaproteobacterial endosymbiont of cicadas, ’*Candidatus* Hodgkinia cicadicola’ (McCutcheon *et al*., 2009) (hereafter *Hodgkinia*), twice in the betaproteobacterial symbionts of different Auchenorrhyncha (Insecta: Hemiptera), ’*Candidatus* Zinderia insecticola’ and ’*Candidatus* Nasuia deltocephalinicola’ (Bennett and Moran, 2013) (hereafter *Zinderia* and *Nasuia*, respectively), and once in the gammaproteobacterial symbionts of Cassidinae beetles, ’*Candidatus* Stammera capleta’ (Salem *et al*., 2017) (hereafter *Stammera*).

**Figure 3.**
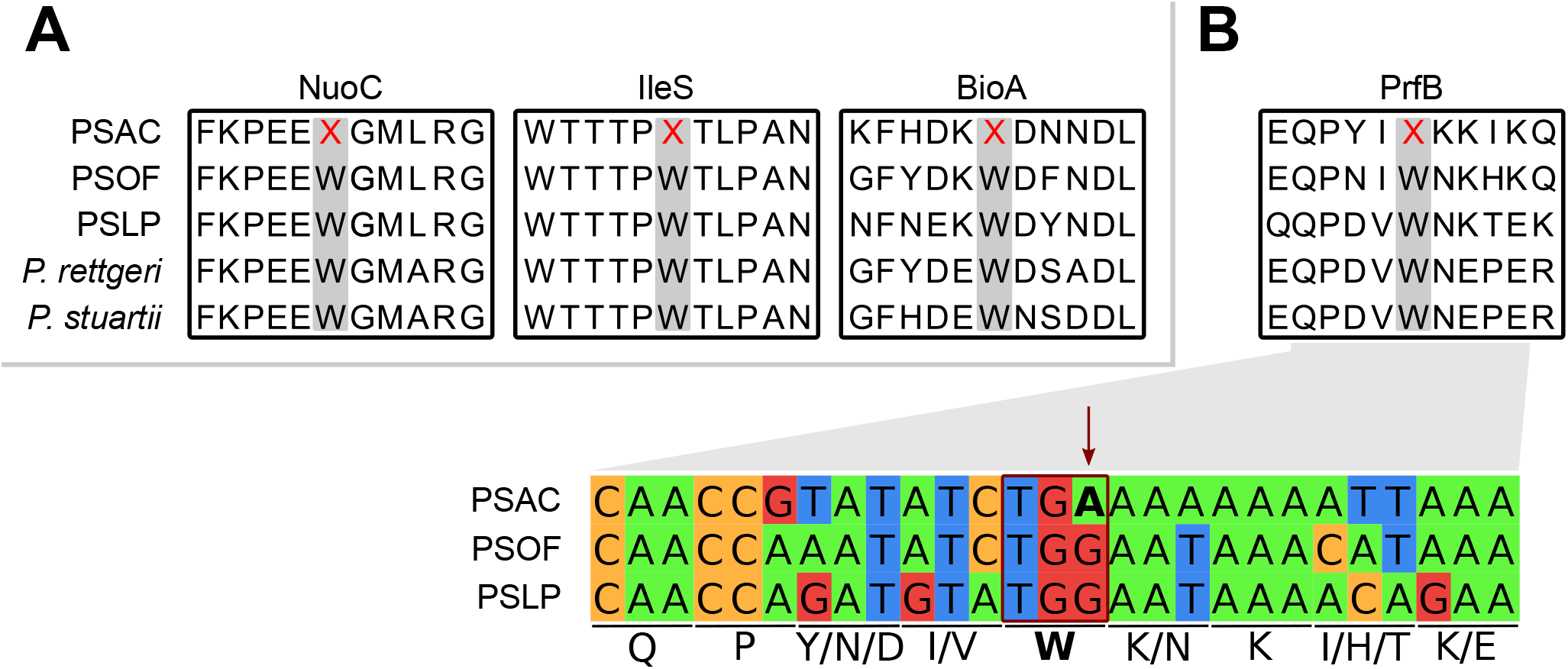
Schematic representation of genetic re-coding in PSAC. **(A)** Excerpt of alignments displaying the translated UG**A** codons found in PSAC compared to PSOF, PSLP, and the free-living *P. rettgeri* strain AR 0082 and *P. struartii* strain MRSN 2154. An “X” in the alignments represents a UG**A** codon. **(B)** Excerpt of alignment displaying the translated UG**A** codon in the same fashion as in A, and including the nucleotide alignment. Under the sequence of each codon, the encoded amino acids are shown with the one-letter abbreviation code.

By comparing PSAC to these other endosymbiont lineages showing genetic re-coding, it becomes apparent that they all show differences in two key factors behind the evolution of the alternative genetic code: the presence/absence of the *prfB* gene and the *trnW*’s **C**CA/**U**CA anticodon, preferentially recognising the codons UG**G** and UG**A**, respectively (**table 2**). Both *Hodgkinia* and *Nasuia* show a loss of the *prfB* gene and the re-coding of the *trnW* anticodon from **C**CA>**U**CA, while *Stammera* and *Zinderia* do not display a change in the *trnW*’s anticodon. Despite this difference, all four endosymbionts have a large proportion of their genes carrying a tryptophan-coding UG**A** codon, from 50.37% in *Nasuia* up to 70.41% for *Hodgkinia*. As for PSAC, it is the only re-coded endosymbiont that retains both a *trnW*-(**C**CA) (with only two unique mutations in the arm loops, supplementary **figure S4**) and a *prfB* gene. However, the *prfB* gene of PSAC itself carries a UG**A** codon (**figure 3B**), which would render it a pseudogene under the genetic code 11 but not under genetic code 4. In addition, only 20.49% of its protein-coding genes carry a UG**A** stop codon, which is in sharp contrast to what is observed in *Stammera, Hodgkinia, Zinderia*, and *Nasuia*. Furthermore, PSAC retains at least nine genes that potentially still use a UG*A* stop codon: *rpsK, fabZ, map, tamA, purE, ispB, rnc, kdsA*, and *miaA*. Without a functioning UG**A** stop codon, the previously mentioned genes would produce rather large proteins towards their 3’-end, putatively rendering them non-functional. Lastly, by re-analysing the annotation of PSOF and PSLP, we also found nine genes in each genome that are predicted to be pseudogenised due to a nonsense mutation (UG**G**>UG**A**; **table 3**). Many of these genes are considered essential in *E. coli* and/or also preserved in other endosymbionts with highly reduced genomes. It is noteworthy the case of *rimM*, which despite conflicting evidence for its essentiality in *E. coli* (judged in EcoCyc (Keseler *et al*., 2017) as ”growth”/”no growth”), it is considered an essential gene, as different mutant-carrying strains exhibit growth defects and a slower rate of translation compared to the wild type. In PSOF and PSLP, *rimM* would be pseudogenised unless translational readthrough occurs to interpret the in-frame UG**A** codon as tryptophan. Lastly, while many of the UG**A**-containing genes in PSOF and PSLP also contain an UG**A** codon coding for tryptophan in PSAC; *apbE, lolA*, and *rimM* share the same tryptophan-encoding UG**A** codon, suggesting an ancestral state for this trait that has been conserved throughout evolutionary time.

**Table 2.**
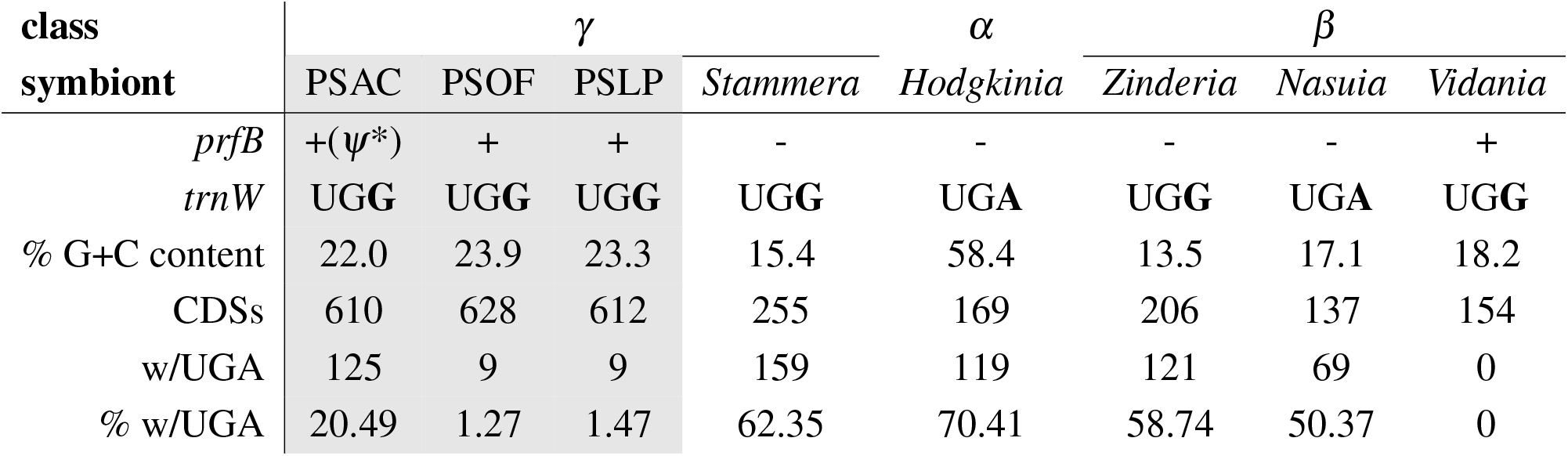
Characteristics accompanying genetic code changes in selected animal endosymbionts. * Indicates the presence of a UG**A** stop codon in the *prfB* pseudogene. Newly sequenced organisms are highlighted in grey background. Accession numbers are found in supplementary **table S2**. *α*=Alphaproteobacteria; *β*=Betaproteobacteria; *γ*=Gammaproteobacteria.

**Table 3.** Genes in PSOF and PSLP predicted produce full-length proteins through translational readthrough of the UGA codon. * Essentiality is denoted in the table as + (yes), -(no), or m(mixed) in the format of +/+/+ (essentiality in *E. coli* as recorded in EcoCyc as ”Growth” or ”No Growth” Keseler *et al*. (2017)/retention in the genomes of *B. aphidicola*, ’*Ca*. Sulcia muelleri’, and ’*Ca*. Blochmannia’ strains/retention in the genome of PSAC).

| gene | Product | Function | Essentiality* | Symbiont |
| --- | --- | --- | --- | --- |
| *apbE* | FAD:protein FMN transferase | Flavin transferase | -/m/+ | PSOF, PSLP |
| *dnaG* | DNA primase | Replication | +/+/+ | PSOF |
| *epmB* | L-lysine 2,3-aminomutase | Isomerase | -/m/+ | PSOF |
| *folA* | Dihydrofolate reductase | Folic acid biosynthesis | +/m/+ | PSLP |
| *ispE* | 4-diphosphocytidyl-2-C-methyl-D-erythritol kinase | Isopentenyl diphosphate | +/m/+ | PSLP |
| *lolA* | Outer-membrane lipoprotein carrier protein | Lipoprotein translocation | m/m/+ | PSOF, PSLP |
| *menC* | *o*-succinylbenzoate synthase | Menaquinone biosynthesis | -/-/+ | PSOF |
| *mepS* | Murein DD-endopeptidase<br>MepS/Murein<br>LD-carboxypeptidase | Cell wall biosynthesis | -/-/+ | PSLP |
| *pdxH* | Pyridoxine/pyridoxamine 5'-phosphate oxidase | PLP biosynthesis | m/m/+ | PSOF |
| *prmC* | Release factor glutamine methyltransferase | Peptide chain release factor methyltransferase | +/m/+ | PSLP |
| *rimM* | Ribosome maturation factor RimM | Ribosome assembly | m/+/+ | PSOF, PSLP |
| *sufS* | Cysteine desulfurase | Iron-sulfur cluster assembly | -/+/+ | PSLP |
| *tilS* | tRNA(Ile)-lysidine synthase | TRNA modification | +/+/+ | PSLP |
| *tsaC* | Threonylcarbamoyl-AMP | TRNA modification | m/m/+ | PSOF |
| *znuB* | High-affinity zinc uptake system membrane protein<br>ZnuB | Zn<sup>2+</sup> uptake | -/m/+ | PSOF |

## Discussion

When endosymbionts experience regular vertical transmission and become locked into the symbiotic association with their hosts, they undergo continuous bottlenecks every new host generation that results in many of the typical genomic characteristics observed in long-term strict vertically-transmitted endosymbionts (Latorre and Manzano-Marín, 2017; Moran *et al*., 2008). These changes typically include a large-scale genome reduction, biased nucleotide composition, a compact gene-dense genome, and a lack of mobile elements. *P. siddallii*, the obligate bacterial endosymbionts of *Haementeria* leeches, display such genomic characteristics, suggesting a long-term association with their hosts coupled with strict vertical transmission. The perfect conservation of B-vitamin biosynthetic genes across *P. siddallii* strains, despite large-scale genome reduction, points towards this symbiotic machinery being key to the establishment and maintenance of this symbiont lineage in *Haementeria* leeches, and provides further evidence that this association is essential for the survival of the leeches. Relating to the loss of *nudB* across *P. siddalli*, it has been shown that, in *Escherichia coli*, many phosphatases show wide-range substrate specificities (Haase *et al*., 2013; Kuznetsova *et al*., 2006), which suggests other phosphatase(s) encoded in the *P. siddalli* genomes might be fulfilling *nudB*’s role. Together, the similarly-reduced genomes, the large shared gene fraction, and the perfect genome synteny across *P. siddallii* strains provide strong evidence for an evolutionary history marked by an early and rapid genome reduction followed by diversification with their leech hosts. This genome reduction, followed by host-symbiont co-divergence and large-scale genome synteny, mirrors well-known examples of ”ancient” nutritional symbionts of aphids (Chong *et al*., 2019; Tamas *et al*., 2002), *Camponotus* ants (Williams and Wernegreen, 2015), cockroaches, and termites (Kinjo *et al*., 2018), among many others.

As we have shown, the genome of PSAC, the obligate nutritional symbiont of the blood-feeding leech *H. acueceyetzin*, does not only show the ”typical” genomic characteristics of a long-term vertically transmitted endosymbiont, but has also evolved a lineage-specific alternative genetic code, reflected in at least 20.49% of its encoded proteins. The alternative genetic code matches the so-called genetic code 4, which features a UG**A** codon reassignment from STOP to tryptophan (Suzuki and Nagao, 2021). Alternatively, it can be proposed that the ancestor of *P. siddallii* used genetic code 4, and this was subsequently reverted to 11. However, this alternative scenario would be less parsimonious, as it requires two steps instead of one (given that *Providencia* and *Enterobacterales* in general use genetic code 11). Also, and to our knowledge, reversal from genetic code 4 to 11 has not been documented, while the change from 11 to 4 has occurred independently in several bacterial lineages. There are two main players involved in such a genetic code reassignment: the peptide release factor 2 (encoded by *prfB*) and the anticodon of the transfer RNA tryptophan (*trnW*). While the former is in charge of recognising the UG**A** codon and, in response, direct the termination of translation (Capecchi, 1967; Capecchi and Klein, 1970; Scolnick *et al*., 1968), the *trnW*-(**C**CA) gene binds the UG**G** codon directing the addition of tryptophan to the nascent chain. It is possible, however, that due to a mistake or the incorporation of a suppressor tRNA (tRNA^Sec^), readthrough of UG**A** stop codon occurs (Tate *et al*., 2001). In fact, it has been shown in *Escherichia coli* that the efficiency of the readthrough of a UG**A** codon by *trnW* is dependent on its 3’-end context, with an “A” residue following the STOP codon permitting more efficiently the readthrough process (Engelberg-Kulka, 1981; Kopelowitz *et al*., 1992). Therefore, the presence of ”A” residues downstream of the *prfB* gene in *P. siddallii*, and markedly in PSAC (seven ”A”), has likely facilitated the maintenance of a UG**A** codon in PSAC that allows for some level of translational readthrough. Thus, we can hypothesise that this readthrough results in functional protein being produced, albeit at a likely reduced level, triggering the start of a genome-wide genetic code change.

The STOP to Tryptophan codon reassignment is known to have arisen in at least four distantly related endosymbiotic lineages: once in the gammaproteobacterial endosymbiont of tortoise beetles (*Stammera*) (Salem *et al*., 2017), once in the alphaproteobacterial endosymbiont of cicadas (*Hodgkinia*) (McCutcheon *et al*., 2009) and twice in the betaproteobacterial endosymbionts of Auchenorrhyncha (*Zinderia* and *Nasuia*) (Bennett and Mao, 2018; Bennett and Moran, 2013). In these endosymbionts, the complete loss of the *prfB* gene is a constant, and this is accompanied by a large amount (*>*50%) of the protein-coding genes that include a tryptophan-encoding UG**A** codon. On the contrary, PSAC’s genome only has around 20% of such proteins, with many being predicted as essential to the symbiont. This is likely to be explained by both the retention of a *trnW*-(**C**CA) and a *prfB* gene, which is in contrast to the afore-mentioned endosymbionts. Despite the retention of a *prfB* gene, it itself has an in-frame UG**A** codon, a feature that likely leads to a situation where translation of *prfB* occurs sub-optimally. This suboptimal translation might have historically been a trigger for the accumulation of in-frame UG**A** codons in essential proteins. In support of this hypothesis, we found that PSAC likely retains several genes that likely still use a UG**A** stop codon, including the gene encoding for the essential ribosomal protein S11 (*rpsK*). In this vein, retention of *bona fide* stop codons that use the now sense codons (under the alternative genetic code), has also been observed in *Blastocrithidia* trypanosomatids (Záhonová *et al*., 2016), despite its molecular basis remaining elusive. The retention of a UG**A** STOP codon in the aforementioned likely essential genes and *prfB*, in addition to the comparatively low percentage of UG**A**-containing genes in PSAC, points to an early stage of genome recoding in this symbiont. Moreover, we found evidence suggesting that the related symbionts PSOF and PSLP likely depend on the read-trough of in-frame UG**A** stop codons for the correct translation of several likely essential proteins. This finding suggests PSOF and PSLP show features of an early on-set of STOP*>*Tryptophan re-coding, while preserving an intact *prfB* gene and a *trnW*-(**C**CA).

Contrary to what has been proposed for *Hodgkinia* (McCutcheon *et al*., 2009), the results of this study show that in *P. siddallii* a UG**G***>*UG**A** mutation in the *prfB* gene, rather than a *trnW* **C**CA*>***U**CA mutation, initiated the genetic recoding and likely occurred in a background that included translational readthrough of certain essential genes (**figure 4**). This pre-existence of UG**A**-containing genes could have facilitated the initial stages of re-coding and is likely the route that has been followed by at least *Zinderia*, where the *trnW* also keeps a **C**CA anticodon. In PSAC, some readthrough of the *prfB* likely occurs, albeit at a low level, to produce functional PrfB protein, which allows some UG**A** to be interpreted as stop codons while also keeping the presence of in-frame UG**A** tryptophan codons relatively low across the genome. The expected and subsequent loss of the *prfB* gene would be facilitated when no UG**A** stop codon is needed to preserve functional essential proteins. Lastly, the **C**CA*>***U**CA mutation in the *trnW* anticodon would eventually arise, facilitating the maintenance and spread of the new genetic code across the protein-coding genes.

**Figure 4.**
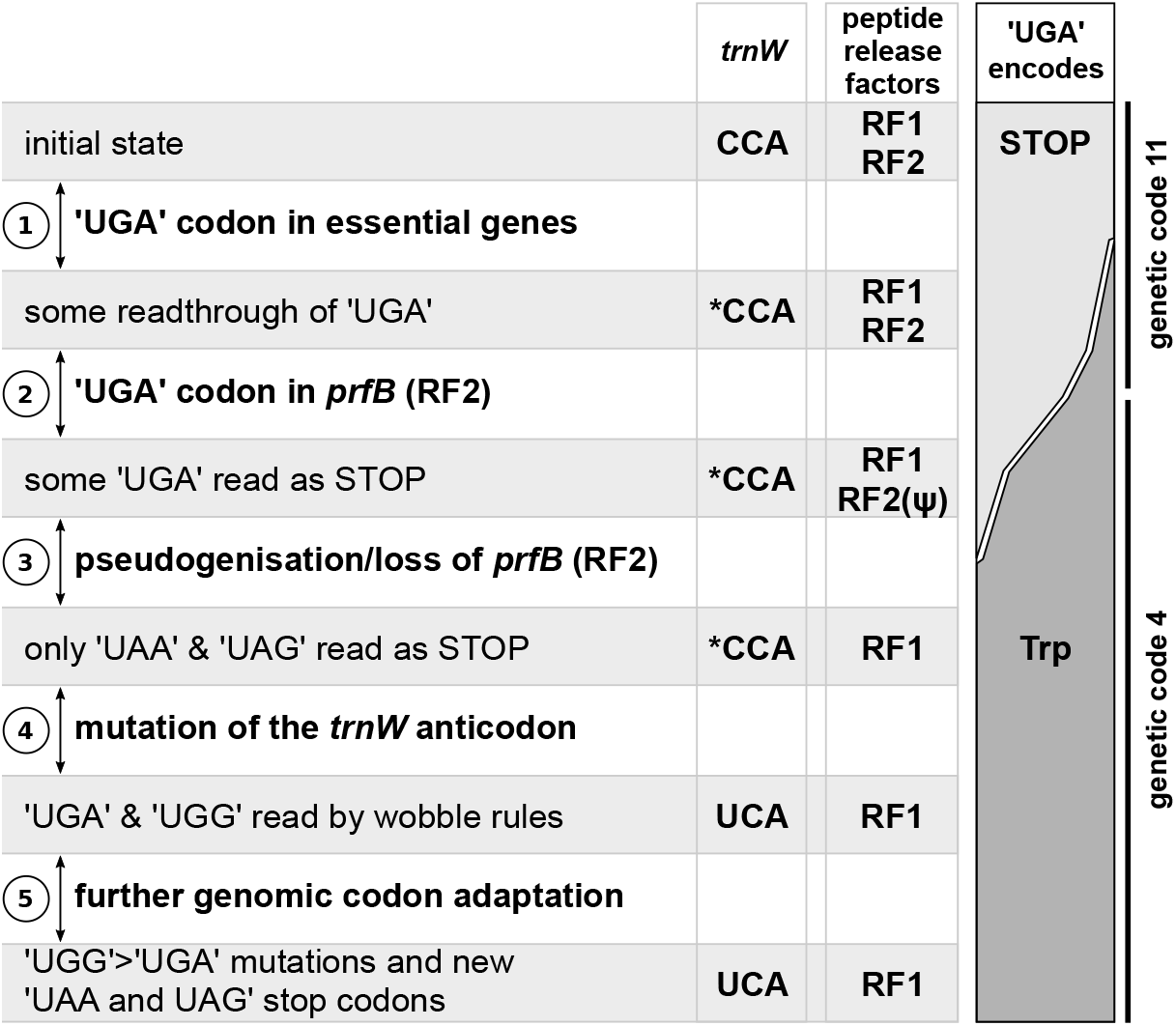
Model showing the step-wise process of ‘UGA’ STOP*>*Trp recoding in PSAC. The asterisk preceding the *trnW* anticodon refers to a tRNA that has undergone distal mutations which allow A-C mismatches at the indicated position. At the right, cartoon diagram displaying the progression of ’UG**A**’ codons being read as STOP/Trp when recoding from gene code 11 to 4. RF1=Peptide chain release factor 1 (*prfA*), RF2=Peptide chain release factor 2 (*prfB*). Figure was based on McCutcheon *et al*. (2009) to facilitate comparison with it.

In conclusion, we found that *P. siddallii* followed a similar genome evolution route to that of *Buchnera aphidicola* and other ”ancient” nutritional endosymbionts, which included a large genome reduction, genome stasis, and loss of mobile elements preceding a co-divergence with their leech hosts. We found conclusive evidence of the lineage-specific evolution of an alternative genetic code in PSAC, in addition to hints of the presence of translational readthrough of some UG**A**-containing genes in the related strains PSOF and PSLP. The findings presented in this study imply that the symbiotic taxon *P. siddallii* is likely one prone to alternative genetic code evolution, having members such as PSOF and PSLP already dependent on some level of translational readthrough. This read-through would thus compensate for nonsense mutations in essential genes to the bacteria or its symbiotic role. We expect further study of the transcriptomics and proteomics of *P. siddallii* strains across the *Haementeria* genus will shed light on the step-wise process of the evolution of an alternative genetic code.

## Materials and Methods

### Leech collection, DNA extraction, and genome sequencing

#### Haementeria acuecueyetzin

A total of 30 *H. acuecueyetin* individuals were collected in Teapa, Tabasco, Mexico in 2018. From these, the 2 pairs of bacteriomes were dissected and DNA was extracted using the *DNEasy Blood & tissue Kit* (Qiagen). DNA library was constructed using the *NGS Nextera Flex DNA library preparation kit* (Illumina). DNA was multiplexed together with 11 other samples and sequenced on a single *HiSeqX* lane (150 base-pairs [**bp**] paired-end reads).

#### Haementeria officinalis

One single individual was collected in Coroneo, Guanajuato, Mexico in 2019. This is the same locality where previous individuals were collected to produce a draft genome of the *P. siddallii* symbiont of *H. officinalis* (Manzano-Marín *et al*., 2015). The bacteriomes were dissected and sorted in absolute ethanol and then used for DNA extraction following a modified CTAB method. Briefly, bacteriomes were frozen in liquid nitrogen and ground with a pestle. 500 μl of CTAB lysis buffer and 10 μl of Proteinase K (20 mg/ml; Promega) were added to the sample and incubated at 65ºC for 90 minutes. Samples were then allowed to cool for 5 minutes and 5μl of RNase (DNase-free; Promega) was added and incubated at 37ºC for 10 minutes. Afterwards, two phenol-chloroform isoamyl alcohol extractions were performed, followed by centrifugation for 10 min at 4ºC after each extraction. Following extractions, 500μl of Chloroform:Isoamyl alcohol (24:1) were added to the resulting upper aqueous phase and centrifuged for 10 minutes at 4ºC. After removing the resulting aqueous phase, 0.1 volumes of ammonium acetate were and 1 volume of isopropanol were added to the sample and incubated at -20ºC overnight. Next, the sample was centrifuged at 14,100 rpm for 15 minutes at 4ºC and supernatant was discarded. Lastly, the DNA pellet was washed twice in 700 μl of 70% ethanol and after drying, it was resuspended in 30 μl of TE 1X buffer. DNA library was constructed using the *Westburg NGS DNA Library Prep Kit* and sequenced in a single run of Illumina *iSeq 100* (150 bp paired-end reads).

#### Haementeria lopezi

One single individual was collected in Cerro colorado, Colima, Mexico in 2022. The bacteriomes were dissected and DNA was extracted using the *High Molecular Weight DNA extraction kit* (Promega) following the plant tissue protocol. DNA library was constructed using the *Westburg NGS DNA Library Prep Kit* and sequenced in a single run of Illumina *MiSeq Micro* (150 bp paired-end reads).

All leech specimens were collected under the permit number 03184 issued by the *Secretaría del Medio Ambiente y Recursos Naturales* (SEMARNAT), to A.O.F.

### Symbiont genome assembly, annotation, and comparative genomics

Reads generated from the Illumina sequencing were right-tail clipped (quality threshold of 20) using FASTX-Toolkit v0.0.13 (http://hannonlab.cshl.edu/fastx_toolkit/, last accessed July 1, 2023). Then, PRINSEQ v0.20.4 (Schmieder and Edwards, 2011) was used to remove reads containing undefined nucleotides (”N”), shorter than 75 bp, and those left without a mate after the trimming and filtering process. The surviving reads were the assembled using SPAdes v3.10.1 (Bankevich *et al*., 2012) with the --only-assembler option and kmers of 77, 99, and 127. The resulting contigs larger than 200 bp were binned using results from a BlastX v2.11.0 (Altschul *et al*., 1997) search (best hit per contig) against a database consisting of the proteome of *Hellobdella robusta* as well as that of a selection of *Providencia* strains (supplementary **table S2**). In all cases, this resulted in one or two large contigs with coverage *>*=20X being assigned to the *Providencia* bin. The completeness of molecules assigned to *Providencia* was corroborated by both manual inspection of contigs showing a higher coverage than the background (*i*.*e*. host contigs) using the BlastX webserver (*vs*. nr) as well as by inspection of assembly graphs. In no case did we find any additional contigs belonging to *Providencia* symbionts. These contigs were used for mapping back the reads from each *Haementeria* species followed by re-assembly of mapped reads using SPAdes as described above. This assembly resulted in a single circular bacterial contig for each *Providencia* endosymbiont.

For creating a draft annotation of the circular chromosomes, we used Prokka v1.14.16 (Seemann, 2014) with a custom protein database built with the predicted proteome of a previously reported draft genome of the *P. siddallii* symbiont of *H. officinalis* (Manzano-Marín *et al*., 2015). This draft annotation was followed by manual curation of the gene coordinates as well as an update to the product naming using UniProtKB (Bateman *et al*., 2021), Pfam v34 (Mistry *et al*., 2021), and InterProScan (Jones *et al*., 2014). Annotations for non-coding RNAs was refined using Infernal v1.1.4 (Nawrocki and Eddy, 2013), with the Rfam database v14.1 (Kalvari *et al*., 2021), tRNAScan-SE v2.0.9 (Chan *et al*., 2021) and ARAGORN v1.2.41 (Laslett, 2004). Pseudogenes were manually identified through on-line BlastX searches of the intergenic regions as well as through BlastP, DELTA-BLAST (Boratyn *et al*., 2012), and domain searches of the predicted open reading frames. Proteins were considered to be putatively functional if all essential domains for the function were found, if a literature search supported the truncated version of the protein as functional in a related organism, or if the predicted protein displayed truncations but retained identifiable domains. Details of the results of these searches are captured in the annotation files. All manual curation was done using UGENE v1.34.0 (Okonechnikov *et al*., 2012).

The predicted proteomes for *P. siddallii* symbionts were clustered into orthologous groups using OrthoMCLv2.0.9 (Chen *et al*., 2007; Li, 2003) followed by a manual check for the split of distant orthologous proteins. These results were used to plot rearrangements using circos v0.69-9 (Krzywinski *et al*., 2009). Single-gene alignments were done using MAFFT v7.453 (--maxiterate 1000 --localpair; Katoh and Standley, 2013) and B vitamin metabolic pathways were plotted following EcoCyc (Keseler *et al*., 2017) and manually drawn using Inkscape v1.1.2 (https://inkscape.org, last accessed July 1, 2023).

### Phylogenetic reconstruction of *Providencia* spp

In order to infer the phylogenetic positioning of the *P. siddalli* clade as well as to illuminate their phylogenetic relationships, we reconstructed a Bayesian phylogenetic tree. For this purpose, we first extracted all ribosomal proteins from selected *Providencia* spp. and *Moellerella wisconsensis* (supplementary **table S1**). Single-gene alignments were done using MAFFT, as described above, followed by removal of divergent and ambiguously aligned blocks with Gblocks v0.91b (Talavera and Castresana, 2007). Next, proteins were concatenated and Bayesian phylogenetic inference was performed in MrBayes v3.2.7 (Ronquist *et al*., 2012) running two independent analyses with four chains each for up to 300,000 generations and checked for convergence (average standard deviation below 0.001) with a burn-in of 25%. Tree was visualised using FigTree v1.4.4 (http://tree.bio.ed.ac.uk/software/figtree/, last accessed July 1, 2023).

All files relating to orthologous protein grouping and phylogenetic analysis can be found at https://doi-.org/10.5281/zenodo.6539517.

## Supplementary Material & Data Availability

Supplementary **figures S1-4** and **tables S1-2** have been included in this submission. All auxiliary files for other analyses as well as the genomes of *P. siddallii* endosymbionts are available online at https://doi.org/10.5281/zenodo.6539517. Newly sequenced and annotated genomes are in the process of being accessioned at the European Nucleotide Archive (ENA).

## Acknowledgements

A.M.M. has received funding from European Union’s *Horizon 2020 Research and Innovation Programme* under the *Marie Sklodowska-Curie* Grant Agreement No. [840270] (LEECHSYMBIO). S.K. was a supported by an *NSERC Discovery Grant*. A.O.F has received funding from the *Programa de Apoyo a Proyectos de Investigación e Innovación Tecnológica-Universidad Nacional Autónoma de México* No. [IN215722]. The authors thank Prof. Dr. Thomas Rattei and his team for maintaining the *Life Science Compute Cluster* (LiSC; https://cube.univie.ac.at/lisc) that was used for computational analyses. The authors would also like to thank SickKids (Toronto, Canada) and the Vienna BioCenter Core Facilities (https://www.viennabiocenter.org/vbcf/; Vienna, Austria). The authors would also like to thank Víctor Sosa Jiménez for assisting in the collection of leech specimens in México. The funders had no role in study design, data collection and analysis, decision to publish, or preparation of the manuscript.

## Notes

https://doi.org/10.5281/zenodo.6539517

### Competing Interest Statement

The authors have declared no competing interest.

### Summary of Updates

Additional phylogenetic analysis of Providencia symbionts using ribosomal proteins, corrections across the manuscript, and and expanded discussion to draw parallels across the tree of life

## References

2010. Genome sequences of the human body louse and its primary endosymbiont provide insights into the permanent parasitic lifestyle. Proceedings of the National Academy of Sciences, 107(27): 12168–12173. URL https://dx.doi.org/10.1073/pnas.1003379107.

Altschul S. F, Madden T. L, Schäffer A. A, Zhang J, Zhang Z, Miller W, and Lipman D. J. 1997. Gapped BLAST and PSI-BLAST: a new generation of protein database search programs. Nucleic Acids Research, 25(17): 3389–3402. URL https://dx.doi.org/10.1093/nar/25.17.3389.

Bankevich A, Nurk S, Antipov D, Gurevich A. A, Dvorkin M, Kulikov A. S, Lesin V. M, Nikolenko S. I, Pham S, Prjibelski A. D, Pyshkin A. V, Sirotkin A. V, Vyahhi N, Tesler G, Alekseyev M. A, and Pevzner P. A. 2012. SPAdes: a new genome assembly algorithm and its applications to single-cell sequencing. Journal of Computational Biology, 19(5): 455–477. URL https://dx.doi.org/10.1089/cmb.2012.0021.

Bateman A, Martin M. J, Orchard S, Magrane M, Agivetova R, Ahmad S, Alpi E, Bowler-Barnett E. H, Britto R, Bursteinas B, Bye-A-Jee H, Coetzee R, Cukura A, Silva A. D, Denny P, Dogan T, Ebenezer T. G, Fan J, Castro L. G, Garmiri P, Georghiou G, Gonzales L, Hatton-Ellis E, Hussein A, Ignatchenko A, Insana G, Ishtiaq R, Jokinen P, Joshi V, Jyothi D, Lock A, Lopez R, Luciani A, Luo J, Lussi Y, MacDougall A, Madeira F, Mahmoudy M, Menchi M, Mishra A, Moulang K, Nightingale A, Oliveira C. S, Pundir S, Qi G, Raj S, Rice D, Lopez M. R, Saidi R, Sampson J, Sawford T, Speretta E, Turner E, Tyagi N, Vasudev P, Volynkin V, Warner K, Watkins X, Zaru R, Zellner H, Bridge A, Poux S, Redaschi N, Aimo L, Argoud-Puy G, Auchincloss A, Axelsen K, Bansal P, Baratin D, Blatter M. C, Bolleman J, Boutet E, Breuza L, Casals-Casas C, de Castro E, Echioukh K. C, Coudert E, Cuche B, Doche M, Dornevil D, Estreicher A, Famiglietti M. L, Feuermann M, Gasteiger E, Gehant S, Gerritsen V, Gos A, Gruaz-Gumowski N, Hinz U, Hulo C, Hyka-Nouspikel N, Jungo F, Keller G, Kerhornou A, Lara V, Le Mercier P, Lieberherr D, Lombardot T, Martin X, Masson P, Morgat A, Neto T. B, Paesano S, Pedruzzi I, Pilbout S, Pourcel L, Pozzato M, Pruess M, Rivoire C, Sigrist C, Sonesson K, Stutz A, Sundaram S, Tognolli M, Verbregue L, Wu C. H, Arighi C. N, Arminski L, Chen C, Chen Y, Garavelli J. S, Huang H, Laiho K, McGarvey P, Natale D. A, Ross K, Vinayaka C. R, Wang Q, Wang Y, Yeh L. S, and Zhang J. 2021. Uniprot: The universal protein knowledgebase in 2021. Nucleic Acids Research, 49(D1): D480–D489. URL https://dx.doi.org/10.1093/nar/gkaa1100.

Baumann P. 2005. Biology of bacteriocyte-associated endosymbionts of plant sap-sucking insects. Annual Review of Microbiology, 59(1): 155–189. URL https://dx.doi.org/10.1146/annurev.micro.59.030804.121041.

Bennett G. M and Mao M. 2018. Comparative genomics of a quadripartite symbiosis in a planthopper host reveals the origins and rearranged nutritional responsibilities of anciently diverged bacterial lineages. Environmental Microbiology, 20(12): 4461–4472. URL https://dx.doi.org/10.1111/1462-2920.14367.

Bennett G. M and Moran N. A. 2013. Small, smaller, smallest: The origins and evolution of ancient dual symbioses in a phloem-feeding insect. Genome Biology and Evolution, 5(9): 1675–1688. URL https://dx.doi.org/10.1093/gbe/evt118.

Bennett G. M and Moran N. A. 2015. Heritable symbiosis: The advantages and perils of an evolutionary rabbit hole. Proceedings of the National Academy of Sciences, 112(33): 10169–10176. URL https://dx.doi.org/10.1073/pnas.1421388112.

Bennett G. M, McCutcheon J. P, McDonald B. R, and Moran N. A. 2016. Lineage-specific patterns of genome deterioration in obligate symbionts of sharpshooter leafhoppers. Genome Biology and Evolution, 8(1): 296–301. URL https://dx.doi.org/10.1093/gbe/evv159.

Boratyn G. M, Schäffer A. A, Agarwala R, Altschul S. F, Lipman D. J, and Madden T. L. 2012. Domain enhanced lookup time accelerated BLAST. Biology Direct, 7(1): 12. URL https://dx.doi.org/10.1186/1745-6150-7-12.

Bové J. M. 1993. Molecular features of Mollicutes. Clinical Infectious Diseases, 17(Supplement 1): S10–S31. URL https://dx.doi.org/10.1093/clinids/17.Supplement_1.S10.

Campbell J. H, O’Donoghue P, Campbell A. G, Schwientek P, Sczyrba A, Woyke T, Söll D, and Podar M. 2013. Uga is an additional glycine codon in uncultured sr1 bacteria from the human microbiota. Proceedings of the National Academy of Sciences, 110(14): 5540–5545. URL https://dx.doi.org/10.1073/pnas.1303090110.

Capecchi M. R. 1967. Polypeptide chain termination in vitro: isolation of a release factor. Proceedings of the National Academy of Sciences, 58(3): 1144–1151. URL https://dx.doi.org/10.1073/pnas.58.3.1144.

Capecchi M. R and Klein H. A. 1970. Release factors mediating termination of complete proteins. Nature, 226(5250): 1029–1033. URL https://dx.doi.org/10.1038/2261029a0.

Chan P. P, Lin B. Y, Mak A. J, and Lowe T. M. 2021. tRNAscan-SE 2.0: improved detection and functional classification of transfer RNA genes. Nucleic Acids Research, 49(16): 9077–9096. URL https://dx.doi.org/10.1093/nar/gkab688.

Chen F, Mackey A. J, Vermunt J. K, and Roos D. S. 2007. Assessing performance of orthology detection strategies applied to eukaryotic genomes. PLoS ONE, 2(4): e383. URL https://dx.doi.org/10.1371/journal.pone.0000383.

Chong R. A, Park H, and Moran N. A. 2019. Genome evolution of the obligate endosymbiont Buchnera aphidicola. Molecular Biology and Evolution, 36(7): 1481–1489. URL https://dx.doi.org/10.1093/molbev/msz082.

Duron O, Morel O, Noël V, Buysse M, Binetruy F, Lancelot R, Loire E, Ménard C, Bouchez O, Vavre F, and Vial L. 2018. Tick-bacteria mutualism depends on B vitamin synthesis pathways. Current Biology, 28(12): 1896–1902.e5. URL https://dx.doi.org/10.1016/j.cub.2018.04.038.

Elzanowski A and Ostell J. 2019. The genetic codes.

Engelberg-Kulka H. 1981. Uga suppression by normal trna^TrP^ in Escherichia coli: codon context effects. Nucleic Acids Research, 9(4): 983–991. URL https://dx.doi.org/10.1093/nar/9.4.983.

Haase I, Sarge S, Illarionov B, Laudert D, Hohmann H.-P, Bacher A, and Fischer M. 2013. Enzymes from the haloacid dehalogenase (had) superfamily catalyse the elusive dephosphorylation step of riboflavin biosynthesis. ChemBioChem, 14(17): 2272–2275. URL https://dx.doi.org/10.1002/cbic.201300544.

Ivanova N. N, Schwientek P, Tripp H. J, Rinke C, Pati A, Huntemann M, Visel A, Woyke T, Kyrpides N. C, and Rubin E. M. 2014. Stop codon reassignments in the wild. Science, 344(6186): 909–913. URL https://dx.doi.org/10.1126/science.1250691.

Jones P, Binns D, Chang H.-Y, Fraser M, Li W, McAnulla C, McWilliam H, Maslen J, Mitchell A, Nuka G, Pesseat S, Quinn A. F, Sangrador-Vegas A, Scheremetjew M, Yong S.-Y, Lopez R, and Hunter S. 2014. InterProScan 5: genome-scale protein function classification. Bioinformatics, 30(9): 1236–1240. URL https://dx.doi.org/10.1093/bioinformatics/btu031.

Kalvari I, Nawrocki E. P, Ontiveros-Palacios N, Argasinska J, Lamkiewicz K, Marz M, Griffiths-Jones S, Toffano-Nioche C, Gautheret D, Weinberg Z, Rivas E, Eddy S. R, Finn R. D, Bateman A, and Petrov A. I. 2021. Rfam 14: Expanded coverage of metagenomic, viral and microRNA families. Nucleic Acids Research, 49(D1): D192–D200. URL https://dx.doi.org/10.1093/nar/gkaa1047.

Katoh K and Standley D. M. 2013. MAFFT multiple sequence alignment software version 7: Improvements in performance and usability. Molecular Biology and Evolution, 30(4): 772–780. URL https://dx.doi.org/10.1093/molbev/mst010.

Keeling P. J and Doolittle W. F. 1996. A non-canonical genetic code in an early diverging eukaryotic lineage. The EMBO Journal, 15(9): 2285–2290. URL https://dx.doi.org/10.1002/j.1460-2075.1996.tb00581.x.

Keseler I. M, Mackie A, Santos-Zavaleta A, Billington R, Bonavides-Martínez C, Caspi R, Fulcher C, Gama-Castro S, Kothari A, Krummenacker M, Latendresse M, Muñiz-Rascado L, Ong Q, Paley S, Peralta-Gil M, Subhraveti P, Velázquez-Ramírez D. A, Weaver D, Collado-Vides J, Paulsen I, and Karp P. D. 2017. The ecocyc database: reflecting new knowledge about Escherichia coli k-12. Nucleic Acids Research, 45(D1): D543–D550. URL https://dx.doi.org/10.1093/nar/gkw1003.

Kikuchi Y and Fukatsu T. 2002. Endosymbiotic bacteria in the esophageal organ of glossiphoniid leeches. Applied and Environmental Microbiology, 68(9): 4637–4641. URL https://dx.doi.org/10.1128/AEM.68.9.4637-4641.2002.

Kinjo Y, Bourguignon T, Tong K. J, Kuwahara H, Lim S. J, Yoon K. B, Shigenobu S, Park Y. C, Nalepa C. A, Hongoh Y, Ohkuma M, Lo N, and Tokuda G. 2018. Parallel and gradual genome erosion in the Blattabacterium endosymbionts of Mastotermes darwiniensis and Cryptocercus wood roaches. Genome Biology and Evolution, 10(6): 1622–1630. URL https://dx.doi.org/10.1093/gbe/evy110.

Kopelowitz J, Hampe C, Goldman R, Reches M, and Engelberg-Kulka H. 1992. Influence of codon context on uga suppression and readthrough. Journal of Molecular Biology, 225(2): 261–269. URL https://dx.doi.org/10.1016/0022-2836(92)90920-F.

Krzywinski M, Schein J, Birol I, Connors J, Gascoyne R, Horsman D, Jones S. J, and Marra M. A. 2009. Circos: an information aesthetic for comparative genomics. Genome research, 19(9): 1639–45. URL https://dx.doi.org/10.1101/gr.092759.109.

Kuznetsova E, Proudfoot M, Gonzalez C. F, Brown G, Omelchenko M. V, Borozan I, Carmel L, Wolf Y. I, Mori H, Savchenko A. V, Arrowsmith C. H, Koonin E. V, Edwards A. M, and Yakunin A. F. 2006. Genome-wide analysis of substrate specificities of the Escherichia coli haloacid dehalogenase-like phosphatase family. Journal of Biological Chemistry, 281(47): 36149–36161. URL https://dx.doi.org/10.1074/jbc.M605449200.

Laslett D. 2004. ARAGORN, a program to detect tRNA genes and tmRNA genes in nucleotide sequences. Nucleic Acids Research, 32(1): 11–16. URL https://dx.doi.org/10.1093/nar/gkh152.

Latorre A and Manzano-Marín A. 2017. Dissecting genome reduction and trait loss in insect endosymbionts. Annals of the New York Academy of Sciences, 1389(1): 52–75. URL https://dx.doi.org/10.1111/nyas.13222.

Lehane M. J. 2005. Managing the blood meal. In The Biology of Blood-Sucking in Insects, pages 84–115. Cambridge University Press, Cambridge. URL https://dx.doi.org/10.1017/CBO9780511610493.007.

Li L. 2003. OrthoMCL: Identification of ortholog groups for eukaryotic genomes. Genome Research, 13(9): 2178–2189. URL https://dx.doi.org/10.1101/gr.1224503.

Manzano-Marín A, Oceguera-Figueroa A, Latorre A, Jiménez-García L. F, and Moya A. 2015. Solving a bloody mess: B-vitamin independent metabolic convergence among gammaproteobacterial obligate endosymbionts from blood-feeding arthropods and the leech Haementeria officinalis. Genome Biology and Evolution, 7(10): 2871–2884. URL https://dx.doi.org/10.1093/gbe/evv188.

McCutcheon J. P and Moran N. A. 2012. Extreme genome reduction in symbiotic bacteria. Nature reviews. Microbiology, 10(1): 13–26. URL https://dx.doi.org/10.1038/nrmicro2670.

McCutcheon J. P, McDonald B. R, and Moran N. A. 2009. Origin of an alternative genetic code in the extremely small and gc-rich genome of a bacterial symbiont. PLoS Genetics, 5(7): e1000565. URL https://dx.doi.org/10.1371/journal.pgen.1000565.

McCutcheon J. P, Boyd B. M, and Dale C. 2019. The life of an insect endosymbiont from the cradle to the grave. Current Biology, 29(11): R485–R495. URL https://dx.doi.org/10.1016/j.cub.2019.03.032.

Mistry J, Chuguransky S, Williams L, Qureshi M, Salazar G. A, Sonnhammer E. L. L, Tosatto S. C. E, Paladin L, Raj S, Richardson L. J, Finn R. D, and Bateman A. 2021. Pfam: The protein families database in 2021. Nucleic Acids Research, 49(D1): D412–D419. URL https://dx.doi.org/10.1093/nar/gkaa913.

Moran N. A. 1996. Accelerated evolution and muller’s rachet in endosymbiotic bacteria. Proceedings of the National Academy of Sciences, 93(7): 2873–2878. URL https://dx.doi.org/10.1073/pnas.93.7.2873.

Moran N. A, McCutcheon J. P, and Nakabachi A. 2008. Genomics and evolution of heritable bacterial symbionts. Annual Review of Genetics, 42(1): 165–190. URL https://dx.doi.org/10.1146/annurev.genet.41.110306.130119.

Nawrocki E. P and Eddy S. R. 2013. Infernal 1.1: 100-fold faster RNA homology searches. Bioinformatics, 29(22): 2933–2935. URL https://dx.doi.org/10.1093/bioinformatics/btt509.

Nikoh N, Hosokawa T, Moriyama M, Oshima K, Hattori M, and Fukatsu T. 2014. Evolutionary origin of insect-Wolbachia nutritional mutualism. Proceedings of the National Academy of Sciences, 111(28): 10257–10262. URL https://dx.doi.org/10.1073/pnas.1409284111.

Nogge G. 1981. Significance of symbionts for the maintenance of an optimal nutritional state for successful reproduction in haematophagous arthropods. Parasitology, 82(4): 101–104.

Nogge G and Gerresheim A. 1982. Experiments on the elimination of symbionts from the tsetse fly, Glossina morsitans morsitans (diptera: Glossinidae), by antibiotics and lysozyme. Journal of Invertebrate Pathology, 40(2): 166–179. URL https://dx.doi.org/10.1016/0022-2011(82)90112-4.

Oceguera-Figueroa A. 2006. A new species of freshwater leech of the genus Haementeria (annelida: Glossiphoniidae) from jalisco state, mexico. Zootaxa, 1110(1): 39. URL https://dx.doi.org/10.11646/zootaxa.1110.1.4.

Oceguera-Figueroa A. 2008. A new glossiphoniid leech from catemaco lake, veracruz, méxico. The Journal of parasitology, 94(2): 375–80. URL https://dx.doi.org/10.1645/GE-1240.1.

Oceguera-Figueroa A. 2012. Molecular phylogeny of the new world bloodfeeding leeches of the genus Haementeria and reconsideration of the biannulate genus Oligobdella. Molecular Phylogenetics and Evolution, 62(1): 508–514. URL https://dx.doi.org/10.1016/j.ympev.2011.10.020.

O’Hara C. M, Brenner F. W, and Miller J. M. 2000. Classification, identification, and clinical significance of Proteus, Providencia, and Morganella. Clinical Microbiology Reviews, 13(4): 534–546. URL https://dx.doi.org/10.1128/cmr.13.4.534.

Okonechnikov K, Golosova O, and Fursov M. 2012. Unipro UGENE: a unified bioinformatics toolkit. Bioinformatics, 28(8): 1166–1167. URL https://dx.doi.org/10.1093/bioinformatics/bts091.

Perkins S. L, Budinoff R. B, and Siddall M. E. 2005. New gammaproteobacteria associated with blood-feeding leeches and a broad phylogenetic analysis of leech endosymbionts. Applied and Environmental Microbiology, 71(9): 5219–5224. URL https://dx.doi.org/10.1128/AEM.71.9.5219-5224.2005.

Ronquist F, Teslenko M, van der Mark P, Ayres D. L, Darling A, Hohna S, Larget B, Liu L, Suchard M. A, and Huelsenbeck J. P. 2012. MrBayes 3.2: Efficient bayesian phylogenetic inference and model choice across a large model space. Systematic Biology, 61(3): 539–542. URL https://dx.doi.org/10.1093/sysbio/sys029.

Salem H, Bauer E, Kirsch R, Berasategui A, Cripps M, Weiss B, Koga R, Fukumori K, Vogel H, Fukatsu T, and Kaltenpoth M. 2017. Drastic genome reduction in an herbivore’s pectinolytic symbiont. Cell, 171(7): 1520–1531.e13. URL https://dx.doi.org/10.1016/j.cell.2017.10.029.

Sawyer R. T. 1986. Feeding and digestive system. In Leech biology and behaviour. Volume II: Feeding, biology ecology and systematics, pages 467–523. Oxford University Press.

Schmieder R and Edwards R. 2011. Quality control and preprocessing of metagenomic datasets. Bioinformatics, 27(6): 863–864. URL https://dx.doi.org/10.1093/bioinformatics/btr026.

Scolnick E, Tompkins R, Caskey T, and Nirenberg M. 1968. Release factors differing in specificity for terminator codons. Proceedings of the National Academy of Sciences, 61(2): 768–774. URL https://dx.doi.org/10.1073/pnas.61.2.768.

Seemann T. 2014. Prokka: rapid prokaryotic genome annotation. Bioinformatics, 30(14): 2068–2069. URL https://dx.doi.org/10.1093/bioinformatics/btu153.

Shulgina Y and Eddy S. R. 2021. A computational screen for alternative genetic codes in over 250,000 genomes. eLife, 10: e71402. URL https://dx.doi.org/10.7554/eLife.71402.

Siddall M. E, Perkins S. L, and Desser S. S. 2004. Leech mycetome endosymbionts are a new lineage of alphaproteobacteria related to the rhizobiaceae. Molecular Phylogenetics and Evolution, 30(1): 178–186. URL https://dx.doi.org/10.1016/S1055-7903(03)00184-2.

Suzuki T and Nagao A. 2021. Genetic code and its variations. In eLS, pages 147–157. Wiley. URL https://dx.doi.org/10.1002/9780470015902.a0029263.

Talavera G and Castresana J. 2007. Improvement of phylogenies after removing divergent and ambiguously aligned blocks from protein sequence alignments. Systematic Biology, 56(4): 564–577. URL https://dx.doi.org/10.1080/10635150701472164.

Tamas I, Klasson L, Canbäck B, Näslund K. A, Eriksson A.-S, Wernegreen J. J, Sandström J. P, Moran N. A, and Andersson S. G. E. 2002. 50 million years of genomic stasis in endosymbiotic bacteria. Science, 296(5577): 2376–2379. URL https://dx.doi.org/10.1126/science.1071278.

Tamas I, Wernegreen J. J, Nystedt B, Kauppinen S. N, Darby A. C, Gomez-Valero L, Lundin D, Poole A. M, and Andersson S. G. E. 2008. Endosymbiont gene functions impaired and rescued by polymerase infidelity at poly(A) tracts. Proceedings of the National Academy of Sciences of the United States of America, 105(39): 14934–9. URL https://dx.doi.org/10.1073/pnas.0806554105.

Tate W. P, Poole E. S, and Mannering S. A. 2001. Protein synthesis termination. In Encyclopedia of life sciences. John Wiley & Sons, Ltd. URL https://dx.doi.org/10.1038/npg.els.0000549.

Tessler M, de Carle D, Voiklis M. L, Gresham O. A, Neumann J. S, Cios S, and Siddall M. E. 2018. Worms that suck: Phylogenetic analysis of hirudinea solidifies the position of acanthobdellida and necessitates the dissolution of rhynchobdellida. Molecular Phylogenetics and Evolution, 127: 129–134. URL https://dx.doi.org/10.1016/j.ympev.2018.05.001.

Trontelj P, Sket B, and Steinbrück G. 1999. Molecular phylogeny of leeches: Congruence of nuclear and mitochondrial rdna data sets and the origin of bloodsucking. Journal of Zoological Systematics and Evolutionary Research, 37(3): 141–147. URL https://dx.doi.org/10.1111/j.1439-0469.1999.00114.x.

Williams L. E and Wernegreen J. J. 2015. Genome evolution in an ancient bacteria-ant symbiosis: parallel gene loss among Blochmannia spanning the origin of the ant tribe camponotini. PeerJ, 3: e881. URL https://dx.doi.org/10.7717/peerj.881.

Yamao F. 1985. Uga is read as tryptophan in Mycoplasma capricolum. Proceedings of the National Academy of Sciences, 82(8): 2306–2309. URL https://dx.doi.org/10.1073/pnas.82.8.2306.

Záhonová K, Kostygov A. Y, Ševćíková T, Yurchenko V, and Eliáš M. 2016. An unprecedented non-canonical nuclear genetic code with all three termination codons reassigned as sense codons. Current Biology, 26(17): 2364–2369. URL https://dx.doi.org/10.1016/j.cub.2016.06.064.

