## Supplementary figures S1-4 for "Evolution of an alternative genetic code in the *Providencia* symbiont of the haematophagous leech *Haementeria acuecueyetzin*"

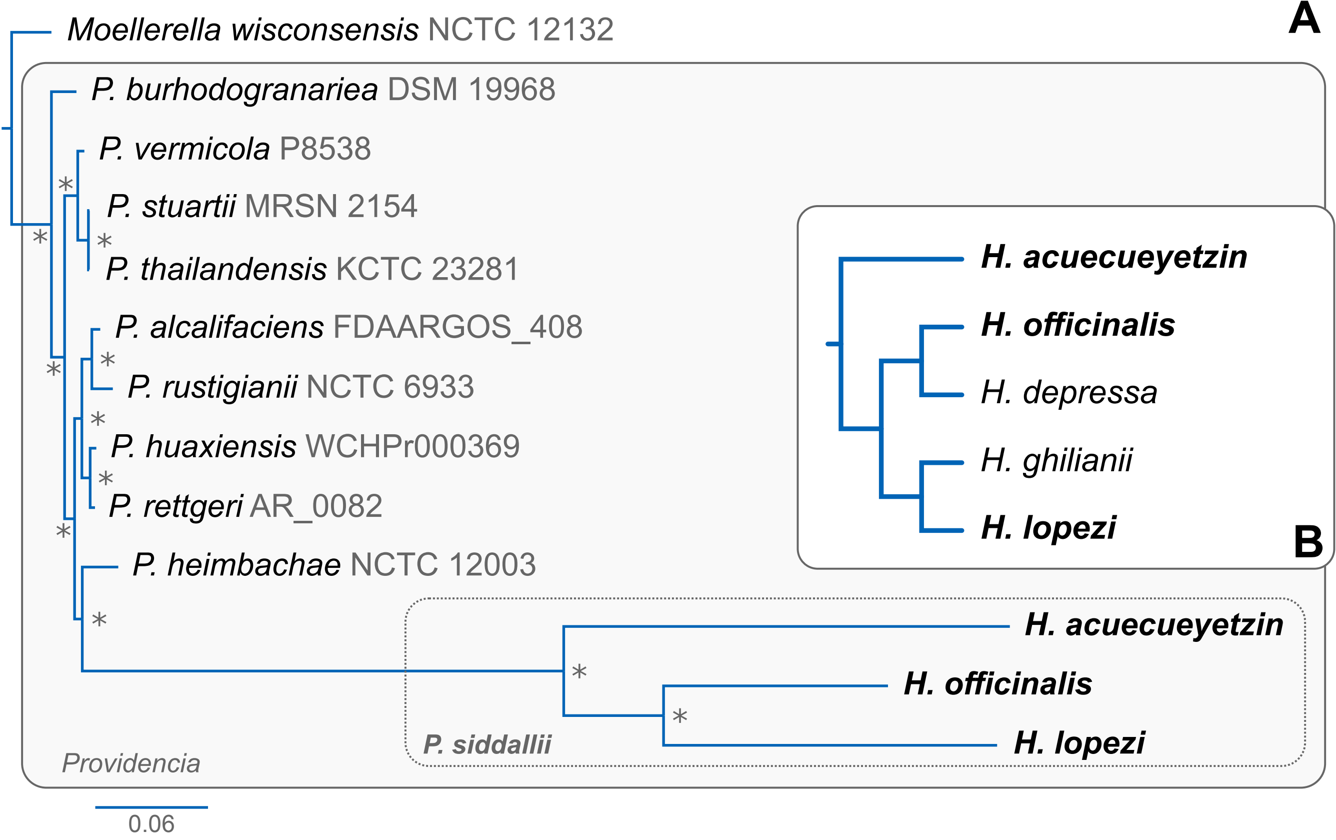


**Figure S1. Phylogenetic relationships of *Providencia* spp. and co-divergence of *Hameneteria-P. siddallii*. (A)** Bayesian phylogenetic tree of *Providencia* spp. based on concatenated ribosomal proteins. Grey boxes delimit the *Providencia* genus- and *P. siddallii* species-level clade. At the leafs, bacterial binomial species are stated with strain designation in dark grey. For the *P. siddallii* clade, the host species name is used instead of the bacterial species and strain. **(B)** Dendrogram displaying the phylogenetic relationships among the *Haementeria* clade of relevance based on **Figure 1**. Species for which endosymbionts were sequenced are highlighted in bold.

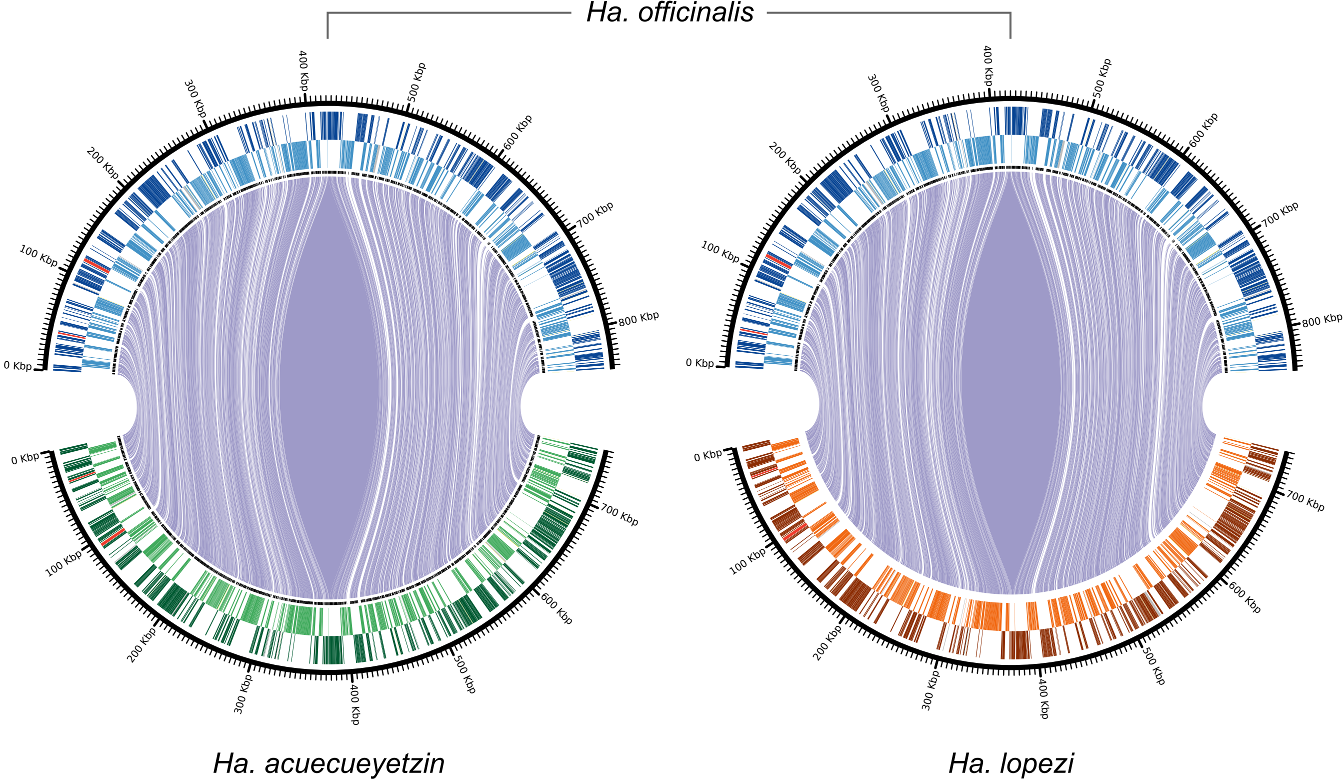


**Figure S2. Conservation of synteny among *P. siddallii* strains.** Circular plots representing linking orthologous proteins between pairs of P. siddallii strains. On the top and bottom of the plots, the leech host name of each endosymbiont is shown. in the outer rims, CDSs coded in the forward strand are shown. In the inner rims, CDSs coded in the reverse strand are shown. Purple lines connect orthologous genes in the same orientation. black bars and red bars represent tRNAs and rRNAs, respectively.

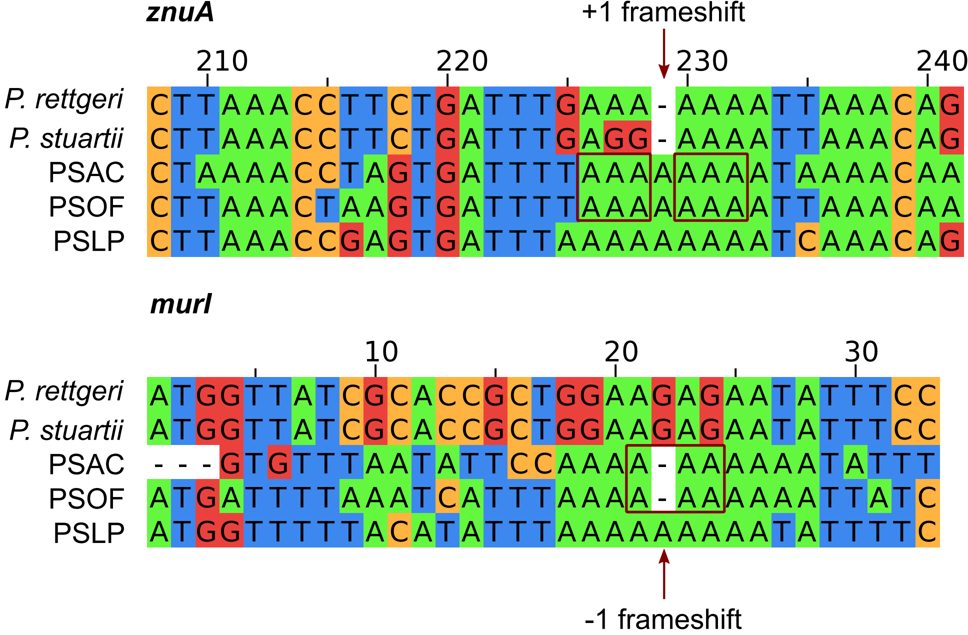


**Figure S3. Predicted transcriptional frameshifted regions in *P. siddallii* genes.** Excerpt of multiple alignments showing examples of a +1 and a -1 frameshift in predicted low-complexity tracks of ***P. siddallii*** genes. Red boxes and arrows showcase the predicted frameshifted region.

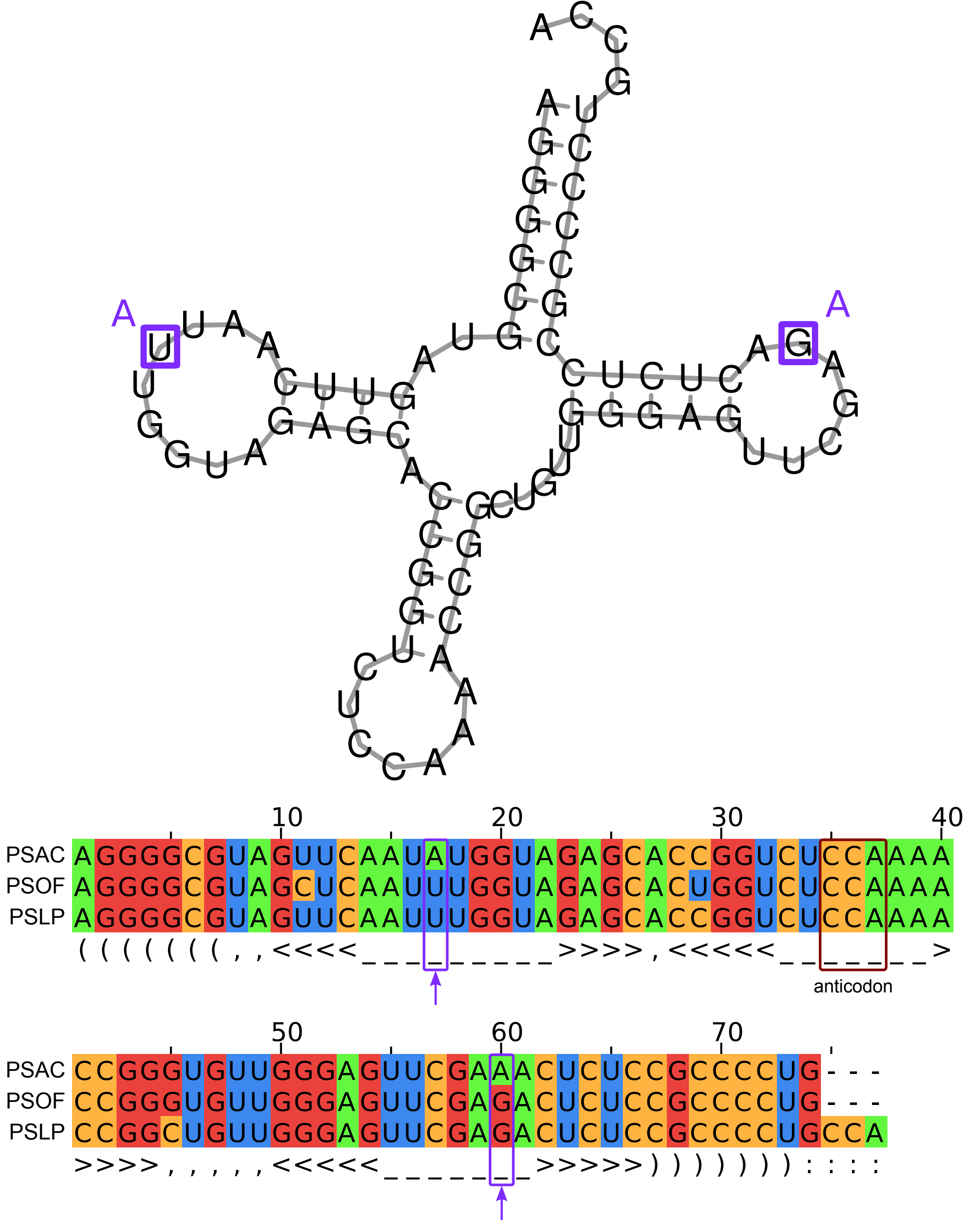


**Figure S4. Structure and mutations in *trnW* genes of *P. siddallii* endosymbionts.** At the top, predicted ***trnW*** RNA structure of PSLP. Residues that show lineage-specific mutations in PSAC are highlighted in light-purple boxes along with the corresponding substitution. At the bottom, structural alignment of ***trnW*** gene sequences of ***P. siddallii*** symbionts. Light-purple and dark-red boxes mark lineage-specific mutations in PSAC and the anticodon, respectively. Under the sequence alignment, the structure of the tRNA is displayed in Stockholm-format, according to the tRNA-scan-SE v2.0.11 bacterial models.
